# Oxaliplatin and 5-Fluorouracil induce p53-p21-pRb-associated cell cycle arrest and a transient senescence-like phenotype in patient-derived low-grade serous ovarian cancer cells

**DOI:** 10.64898/2026.09.22.753519

**Authors:** Rewati Prakash, Benjamin N Forgie, Alicia A Goyeneche, Edith Zorychta, Abu Shadat M Noman, Lucy Gilbert, Carlos M Telleria

## Abstract

**Objectives:** Low-grade serous ovarian cancer (LGSOC) is characterized by frequent *TP53* wild-type status and limited responsiveness to conventional chemotherapy. This study investigated the cytotoxic effects of oxaliplatin (OXP) combined with 5-fluorouracil (5-FU) on patient-derived LGSOC cells.

**Results:** Clinically relevant concentrations of OXP and 5-FU had minimal effects on LGSOC cell viability but markedly reduced cellular proliferation and clonogenic recovery. Combined treatment induced pronounced accumulation of cells in the G0/G1 phase of the cell cycle and increased expression of p53 and p21^Cip1^, accompanied by reduced pRb phosphorylation, consistent with activation of the p53–p21–pRb pathway. OXP+5-FU also increased senescence-associated β-galactosidase (SA-β-Gal) activity, cellular enlargement and flattening, and reactive oxygen species (ROS) production, supporting a senescence-like phenotype. However, after drug withdrawal, treated cells progressively regained proliferative capacity, indicating that the senescence phenotype was transient rather than fully irreversible. Together, these findings show that OXP+5-FU primarily arrests rather than kills LGSOC cells and induces a transient senescence-like phenotype and G0/G1-phase arrest by activating the p53–p21–pRb pathway.

## Introduction

Low-grade serous ovarian cancer (LGSOC) is a rare histological subtype, accounting for approximately 2% of epithelial ovarian cancers and 4.7% of serous ovarian cancers [1]. Unlike the most frequent high-grade serous ovarian carcinoma (HGSOC), LGSOC is predominantly *TP53* wild-type and genomically relatively stable, with recurrent MAPK pathway alterations, including KRAS (16–44%), BRAF (2–20%), and NRAS (∼26%) [2].

Although LGSOC generally follows an indolent course, with 5-year survival of approximately 60–75%, most patients eventually relapse and have limited therapeutic options. Standard treatment consists of cytoreductive surgery followed by platinum-based chemotherapy; however, LGSOC is relatively chemoresistant, with platinum response rates reported as low as 4%. Optimal cytoreductive surgery remains the principal approach associated with prolonged survival [3]. For recurrent KRAS-mutated LGSOC, the RAF/MEK inhibitor avutometinib combined with the focal adhesion kinase (FAK) inhibitor defactinib has recently received FDA approval [4-7] but additional therapeutic strategies are needed.

Oxaliplatin (OXP) and 5-fluorouracil (5-FU) are established treatments for gastrointestinal cancers and have also been investigated for ovarian cancer [8,9]. OXP, a third-generation platinum compound, is approved for metastatic colorectal cancer (CRC) when combined with 5-FU [10]. OXP induces DNA damage and can affect mitochondrial and endoplasmic reticulum function [11], whereas 5-FU inhibits thymidylate synthase and disrupts nucleic acid metabolism by incorporating its metabolites into DNA and RNA [12,13].

Greater sensitivity to OXP and/or 5-FU has been reported in *TP53* wild-type tumors, whereas *TP53* mutations have been linked to reduced sensitivity in experimental models [14-16]. Given the predominance of *TP53* wild-type tumors in LGSOC [17], this provides a biological rationale for investigating OXP+5-FU in this disease. LGSOC and CRC also share frequent RAS–MAPK pathway activation and transcoelomic dissemination within the peritoneal microenvironment [18-20], further supporting the repurposing of this established CRC regimen. Drug repurposing can accelerate therapeutic development by leveraging agents with established pharmacological and clinical profiles [21]. Accordingly, we investigated the effects of combined OXP+5-FU on the growth of patient-derived LGSOC cells.

## Material and Methods

### Cell lines, culture conditions, and treatments

LGSOC cell lines VOA6406, VOA1056, and VOA7681 were provided by Dr. Mark Carey (University of British Columbia, Canada). Their derivation from LGSOC patients has been previously reported (Table S1). We selected the cell lines based on confirmed LGSOC origin, characteristic LGSOC driver mutations, and distinct prior treatment histories. Cells were cultured in a 1:1 mixture of MCDB105 (Sigma-Aldrich, St. Louis, MO, USA) and 199 medium (Thermo Fisher, Waltham, MA, USA), supplemented with 10% FBS (Corning Inc., Corning, NY, USA) and penicillin/streptomycin (Mediatech, Manassas, VA, USA), at 37 °C in 5% CO□.

OXP (Sigma) was dissolved in 5% glucose at 0.25 mg/mL and stored at 4 °C for up to 2 months [22]. Cells were treated with 5.0 µM OXP, corresponding to the reported maximum plasma concentration (Cmax) and a representative upper concentration intended to minimize off-target effects [23]. 5-FU (Sigma) was dissolved in DMSO at 10 mg/mL and stored at 4 °C for up to 3 months, with a final DMSO concentration of ≤0.1%. We established cell-line-specific 5-FU concentrations using dose-response clonogenic recovery assays and nonlinear regression of survival data (Figure S1): 5.0 µM for VOA1056, 2.0 µM for VOA6406, and 1.5 µM for VOA7681. Cells were treated with OXP or vehicle for 3 h, followed by 5-FU or DMSO vehicle for 72 h.

### Cell viability and proliferation

Cells were seeded in 6-well plates and allowed to adhere for 24 h before treatment. After treatment, we determined viable and total cell numbers using the Guava ViaCount™ reagent (Cytek Biosciences, Fremont, CA, USA) and a Muse® micro-capillary cytometer (Cytek), which distinguishes viable from nonviable cells by differential permeability to DNA-binding dyes.

### Clonogenic survival

To assess residual drug toxicity, cells were treated with OXP+5-FU for 72 h, after which 1,000 viable cells were replated in drug-free medium and cultured for up to 2 weeks. We counted colonies containing ≥50 cells after fixation with 4% paraformaldehyde and crystal violet staining using an inverted light microscope.

### Cell cycle

After 72 h of OXP+5-FU treatment, cells were fixed with 1% paraformaldehyde and stained with propidium iodide (PI) in the presence of RNase A and Triton X-100. DNA content was analyzed on the Muse micro-capillary cytometer to determine the proportions of cells in the G0/G1, S, and G2/M + >4n phases.

### Western blotting

After 72 h of OXP+5-FU treatment, we prepared whole-cell extracts and performed protein quantification, SDS-PAGE, and transfer to PVDF membranes as previously described [24]. We probed membranes with antibodies against β-actin, p53, p21^cip1^, p16^INK4A^, and phospho-pRb. We used appropriate HRP-conjugated secondary antibodies for detection.

### Senescence-associated β-galactosidase staining

Cells were treated with OXP+5-FU for 72 h and analyzed using the Senescence β-Galactosidase (SA-β-Gal) Staining Kit (Cell Signaling Technology, Danvers, MA, USA). Cells were fixed and incubated with an X-gal-based staining solution at pH 6.0 for up to 18 h. We identified senescence-associated β-galactosidase (SA-β-Gal)-positive cells by blue cytoplasmic staining and quantified them as a percentage of total cells using ImageJ (National Institutes of Health, Bethesda, MD, USA). Five fields per well and three wells per treatment group were analyzed. Untreated and etoposide-treated MCF-7 breast cancer cells served as negative and positive controls, respectively.

### Drug withdrawal and cell growth recovery

To assess the persistence and reversibility of the treatment response, we collected cells after OXP+5-FU exposure and replated them in drug-free medium. Cell viability and number were measured every 48 h using the Guava ViaCount™ and the Muse micro-capillary cytometer.

### Oxidative stress

We measured reactive oxygen species (ROS) production after OXP+5-FU treatment using the Muse Oxidative Stress Assay (Cytek) per the manufacturer’s protocol. Cells were incubated with the assay working solution for 30 min at 37 °C, then analyzed on a Muse micro-capillary cytometer. We detected ROS with dihydroethidium (DHE), which produces red fluorescence upon interaction with superoxide and DNA.

### Statistical analysis

We repeated Western blotting, SA-β-Gal, oxidative stress, and drug withdrawal/recovery experiments at least twice and obtained similar results; we performed all experiments in triplicate. We analyzed data using GraphPad Prism (Dotmatics, Boston, MA, USA), expressed as mean ± SD, and considered results significant at p < 0.05. We compared groups using one-way ANOVA with Dunnett’s multiple-comparison test. Significance was indicated as *p < 0.05, **p < 0.01, and ***p < 0.001 versus vehicle (VEH), and #p < 0.05, ##p < 0.01, and ###p < 0.001 versus OXP+5-FU.

## Results

### OXP and 5-FU treatment of LGSOC cell lines has a strong impact on proliferation capability but not viability

LGSOC cells were treated with OXP and 5-FU, both independently and in combination. Despite the known efficacy of OXP and 5-FU in treating CRC, these treatments did not affect viability in LGSOC cells, regardless of cell line, compared with vehicle-treated cells (Figure 1A). Despite the lack of cell death, the OXP+5-FU combination significantly impaired cell proliferation (Figure 1B). This cytostatic effect was demonstrated in a colony formation assay using a low number of pre-treated cells that received no further drug treatment; the treated cells struggled to form colonies (Figure 1C). Interestingly, both drugs affected each cell line differently, but the combination markedly impaired proliferation.

**Figure 1.**
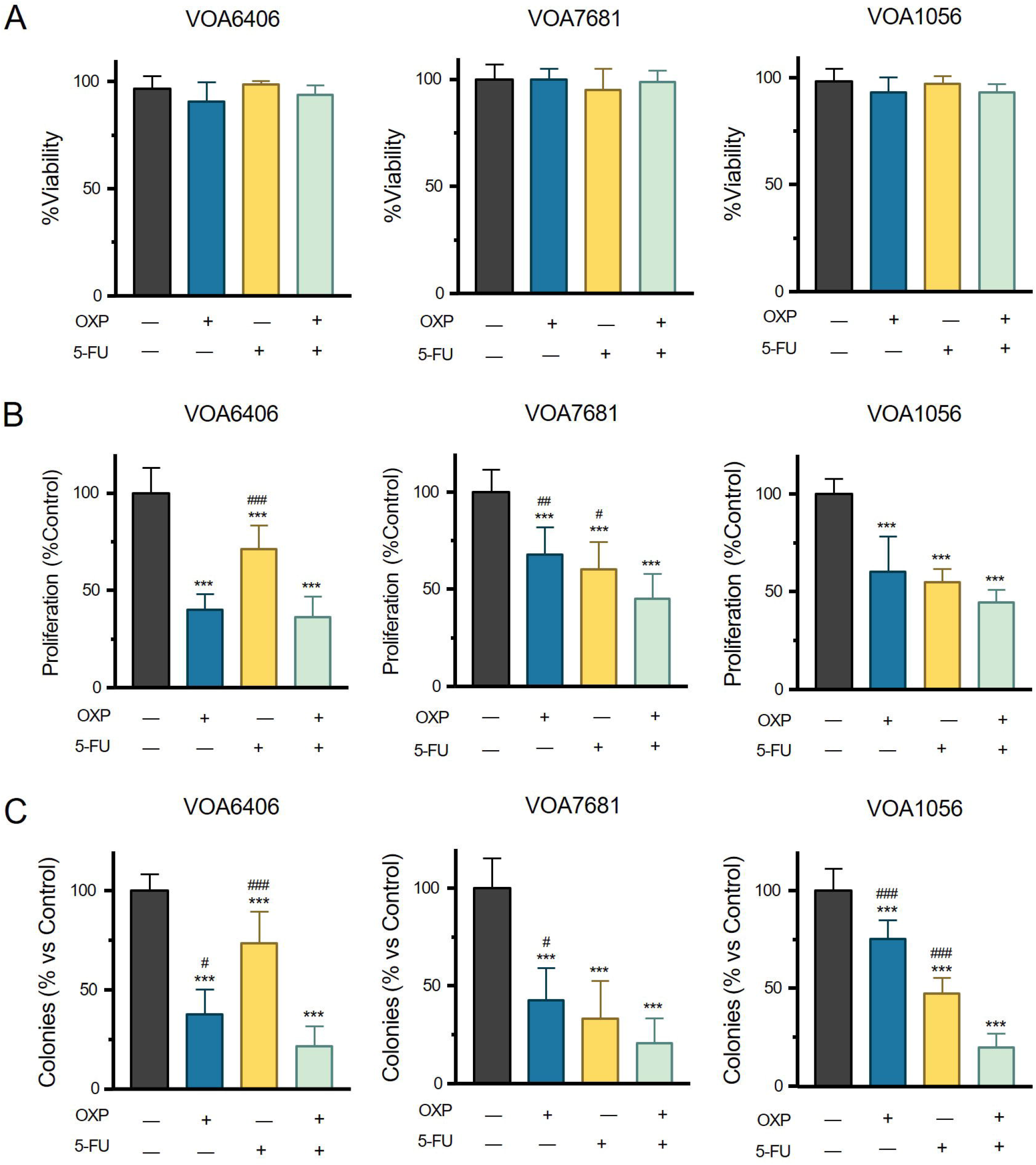
Effect of oxaliplatin (OXP) and 5-fluorouracil (5-FU) on the viability and proliferation of LGSOC cells. After treatment, we assessed viability **(A)** and proliferation **(B).** Cells were then plated for clonogenic recovery **(C)**.

### Combined OXP and 5-FU treatment favors G0/G1 phase cell cycle arrest associated with activation of the p53-p21-pRb axis

Given the established correlation between cytostatic drug effects and cell cycle arrest, our assessment of cell cycle progression revealed a predominant G0/G1 phase accumulation in the LGSOC cell lines treated with both OXP and 5-FU. VOA6406 cells showed G0/G1 rates of 22.78 ± 5.23% (p<0.001) (OXP) and 52.62 ± 8.42% (p<0.001) (OXP+5-FU), respectively, versus 39.82 ± 6.62% (VEH) (Figure 2A). 5-FU-treated VOA6406 cells did not show a significant increase in the percentage of cells in the G0/G1 phase compared with VEH-treated cells. VOA7681 cells showed a similar trend with the drug combination. The G0/G1 populations increased to 76.46 ± 25.77% (p < 0.001) (5-FU) and 83.20 ± 4.10% (p < 0.001) (OXP+5-FU), compared with 64.89 ± 4.59% in VEH-treated cells (Figure 2B). OXP-treated VOA7681 cells showed a negligible reduction in the percentage of cells transiting the G0/G1 phase compared with VEH-treated cells.

**Figure 2.**
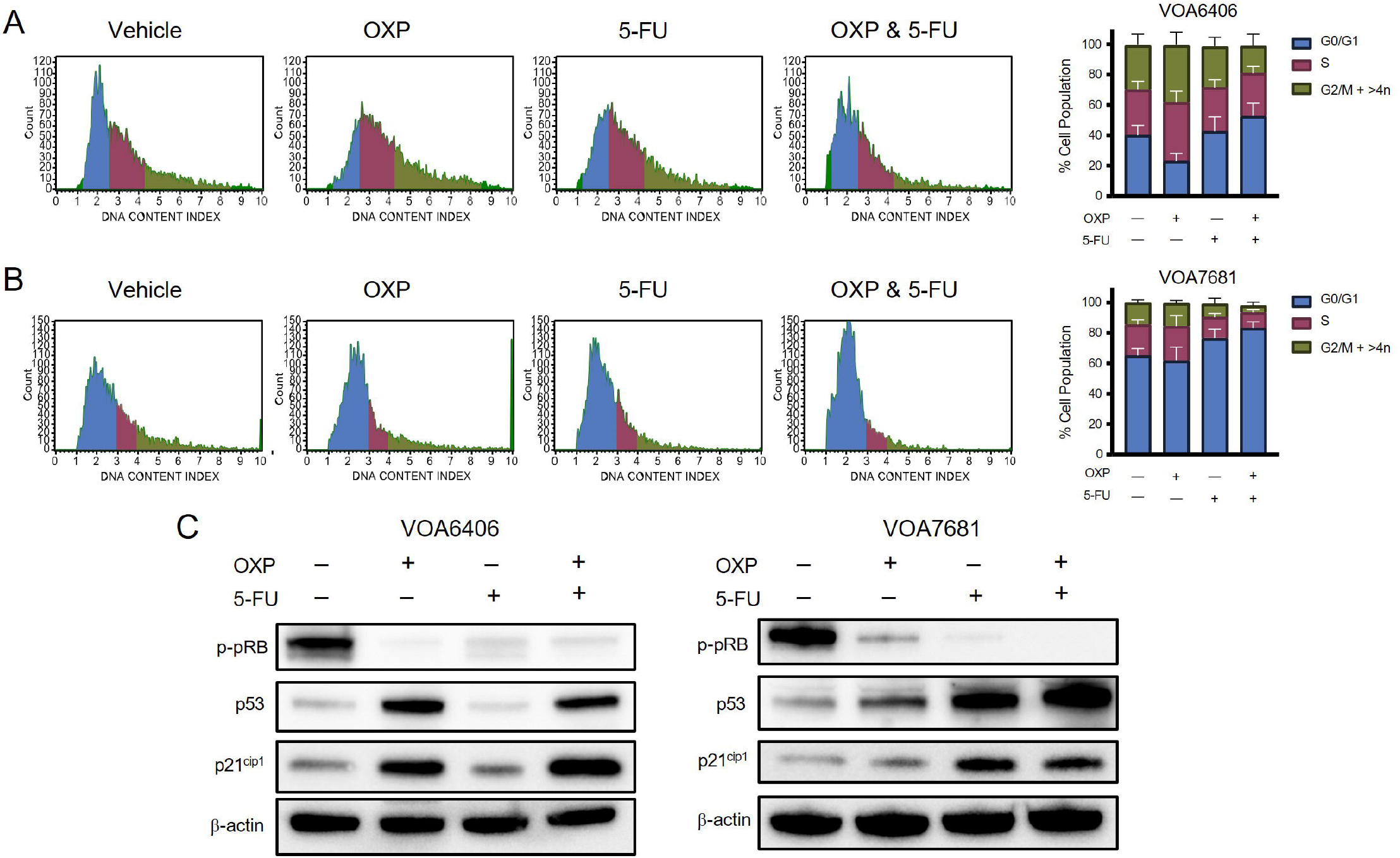
Cell cycle-arresting effects of oxaliplatin (OXP) and 5-fluorouracil (5-FU) in LGSOC cells. Representative flow cytometry histograms showing cell cycle phase populations in VOA6406 **(A)** and VOA7681 **(B).** Effects of OXP, 5-FU, or their combination on retinoblastoma phosphorylation (phospho-pRb) and on the expression of the tumor suppressor protein p53 and the cyclin-dependent kinase inhibitor p21^cip1^, as detected by western blotting. β-actin served as a protein loading control **(C)**.

To further reinforce that OXP+5-FU treatment of LGSOC cells favors arrest in the G0/G1 phase of the cell cycle, we analyzed changes in protein expression associated with arrest at the G1-to-S phase transition. OXP+5-FU distinctly upregulated the tumor suppressor protein p53 and the cyclin-dependent kinase inhibitor (CDKi) p21^cip1^ and promoted dephosphorylation of retinoblastoma protein (pRb) (Figure 2C). Both drugs independently affected protein expression; OXP had a dominant effect in VOA6406, whereas 5-FU was predominant in VOA7681. Notably, in VOA7681, OXP+5-FU further induced p53 expression than either drug alone, an effect not observed in VOA6406.

### Combination of OXP with 5-FU induces a transient senescent-like phenotype in LGSOC cells

Having shown that LGSOC cells treated with OXP+5-FU favor accumulation of cells transiting the G0/G1 phase of the cell cycle with no reduction in viability, we explored the potential for inducing cell cycle arrest-associated cellular senescence. A canonical marker of cellular senescence phenotype in vitro is the activation of the lysosomal protein SA-β-Gal (measured at pH=6.0) [25]. In all three LGSOC cell populations studied, SA-β-Gal staining showed elevated expression in the treatment groups, predominantly in the OXP+5-FU group (Figure 3A). Moreover, positively stained cells also showed increased size and a flattened shape, another common hallmark of cellular senescence (Figure 3B). These images also show that OXP+5-FU-treated cells, particularly in VOA6406, become increasingly granular compared with vehicle-treated cells, indicating likely increased lysosomal activity and cell size. Metabolic changes and adaptations occur during cellular senescence, including ROS production [25]. Measurement of superoxide radical production in VOA6406 and VOA7681 cells showed that OXP+5-FU significantly increases ROS production compared with the other treated groups (Figure 3C).

**Figure 3.**
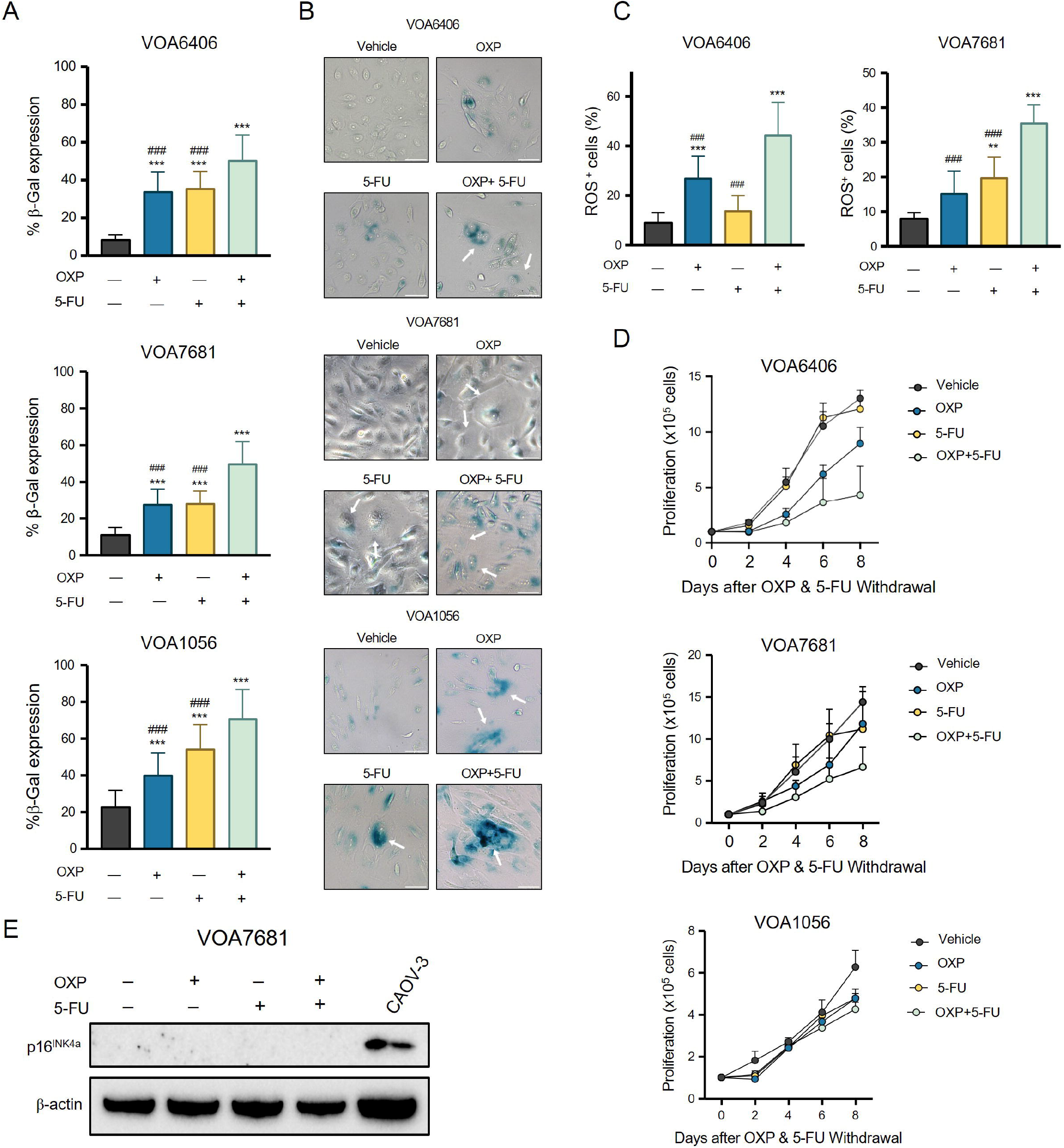
Transient senescent-like phenotype of oxaliplatin (OXP)- and 5-fluorouracil (5-FU)-treated LGSOC cells. Quantification of the SA-β-Gal colorimetric assay **(A)** and representative images of all treatment groups **(B).** We considered blue perinuclear staining a positive result for SA-β-Gal expression. White arrows indicate cellular expansion and flattening. Scale bar=75 µm. Total reactive oxygen species (ROS) production **(C)**. Drug withdrawal and recovery of proliferation in previously OXP+5-FU-treated cells **(D)**. Protein expression of p16^INK4^ in VOA7681 cells, with CAOV-3 cells used as a positive control **(E)**.

To determine whether OXP+5-FU treatment causes irreversible or reversible damage, we used a drug withdrawal and recovery assay (Figure 3D). Notably, all patient-derived LGSOC cell lines eventually returned to normal proliferation, leaving their cell cycle arrest and transient senescent phenotype. In particular, proliferation in the OXP-alone and OXP/5-FU treatment groups remained relatively slow for 2-4 days post-treatment before the cultures began proliferating at a rate similar to the VEH group. However, in VOA6406 and VOA7681, OXP+5-FU-treated cells did not reach 50% of the growth of the vehicle groups by day 8, indicating a small, prolonged toxic effect that extended beyond drug withdrawal.

Alongside activation of the p53-p21^cip1^-pRB axis [26] (Figure 2C), cell cycle transition toward G0/G1 and induction of transient cellular senescence are associated with CDKi activation (e.g., p21^cip1)^ [27] (Figure 2C). Most of the LGSOC cell lines generated in Dr. Carey’s lab harbor loss of the CDKi CDKN2A, resulting in absent or markedly reduced p16^INK4A^ expression (12/14; 86%) [28]. Coincidentally, VOA6406 (Figure 3E), as well as VOA1056 and VOA7681 (data not shown), do not express the p16^INK4^ protein when compared with protein extracts from high-grade serous CAOV-3 ovarian cancer cells (Figure 3E) [29] used as a positive control for p16^INK4^ expression [30].

## Discussion

The OXP+5-FU combination is primarily used for CRC and mucinous ovarian cancer [8,11]. We investigated it in LGSOC because OXP has a bulky carrier ligand that confers a resistance profile distinct from carboplatin, potentially enabling activity in platinum-resistant LGSOC cells [11]. In addition, LGSOC is predominantly *TP53* wild-type, with pathogenic *TP53* mutations reported in only ∼8% of cases. OXP/5-FU responses have been associated with *TP53* wild-type status [16,31,32].

Unlike CRC cells, where clinically achievable OXP/5-FU concentrations can induce cell death, our treatment predominantly inhibited LGSOC proliferation without substantial cytotoxicity. This was associated with transient G0/G1 arrest and p53 activation, which transcriptionally upregulates the CDKi CDKN1A coding for p21^cip1^ [33]. p21^cip1^ promotes inhibition of cyclin-dependent kinases (CDKs) and hypophosphorylation of pRb, maintaining pRb activity and restricting E2F-dependent transcription required for G1-to-S progression [34-36]. Consistent with our findings, OXP+5-FU induces G0/G1 arrest in *TP53* wild-type CRC cells [37,38].

Because OXP+5-FU primarily inhibited proliferation, we investigated whether this response reflected a senescence-like phenotype. Cellular senescence is a stress-responsive state characterized by cell-cycle arrest and accompanied by morphological, metabolic, and secretory changes. Oxidative stress can promote senescence through DNA damage and activation of the p53-p21 and/or p16-pRb pathways [39]. Senescent cells may undergo metabolic remodeling and acquire a senescent-associated secretory phenotype (SASP) [25] .

Our LGSOC models lacked detectable p16^INK4a^ [25,35], which may contribute to the transient response because p16^INK4a^ signaling supports durable senescence [40]. Thus, our findings are more consistent with a transient, reversible senescence-like state than with fully established senescence. González-Gualda *et al*. proposed that senescence should include cell-cycle arrest, a structural change, and an additional phenotype associated with the senescence subtype [25]. OXP+5-FU-treated cells exhibited G0/G1 arrest, cellular flattening, increased SA-β-Gal activity, and oxidative stress, supporting a senescence-like phenotype. However, additional canonical markers and functional assays are needed to establish bona fide senescence.

LGSOC frequently harbors RAS-RAF-MEK-MAPK alterations linked to oxidative stress and senescence-associated responses [41,42]. ROS and p21^cip1^ signaling can reinforce stress-induced cell-cycle arrest [43]. Chemotherapeutic agents can also induce reversible senescence-like states; etoposide and doxorubicin produced such features in non-small-cell lung, colon, and breast cancer cells, with proliferative recovery in some cells as early as day 5 [44]. Consistent with distinct roles in senescence initiation and maintenance, p21^cip1^ can initiate senescence, whereas p16^INK4a^ becomes increasingly important for its maintenance [45]. The absence of p16^INK4a^ in our models [46,47] may therefore permit cell-cycle re-entry after transient p21^cip1^-associated arrest.

This phenotype may represent a therapeutic vulnerability. Senescence-like states can create a window for senolytic therapy, which targets molecular dependencies that support senescent-cell survival [48-50]. For instance, cisplatin induces senescence in *TP53* wild-type and mutant tumor models, and subsequent navitoclax treatment, which inhibits anti-apoptotic Bcl-2 family proteins [51], increases cell death compared with cisplatin alone [52]. These findings suggest a potential sequential “one-two punch”, in which OXP+5-FU induces a reversible senescence-like state, followed by senolytic elimination of surviving cells. This strategy requires validation in LGSOC models.

### Limitations

Although we used clinically achievable concentrations of OXP and 5-FU, differences in cellular transport, protein binding, and cell-cell communication limit extrapolation from these in vitro models. Future studies should therefore evaluate responses in 3D models, including organoids, and in vivo.

The mechanisms underlying the transient phenotype remain incompletely defined. Future studies should determine whether reducing treatment-induced ROS permits cell-cycle re-entry and identify sources of oxidative stress, including mitochondrial dysfunction. The SASP [53] should also be examined; it can influence the tumor microenvironment and chemotherapy response [49].

Caveolin-1, which regulates cell signaling, oxidative stress, and senescence, and is linked to ROS production and p53 stabilization [41,54] may contribute to this phenotype. Its role should be assessed after antioxidant treatment and genetic downregulation to determine whether the phenotype can be rescued [55].

Finally, we did not directly examine the RAS/RAF-MAPK/MEK pathway; oxidative stress can activate p38MAPK, which in turn can activate the p53-p21^cip1^ pathway [41]. Because LGSOC has mutations in the RAS/RAF genes, this pathway is very likely involved through oxidative stress-induced senescence (OIS). Therefore, measuring ERK activity and the mitogen-activated kinases (MKK3 and MKK6) could help elucidate the overall mechanism.

## Supporting information

Uncropped Western Blot Images

## Declarations

### Ethics approval and consent to participate

Not applicable

### Consent for publication

All authors have approved this submission and publication

### Data availability

All data presented are available upon reasonable request to the corresponding author.

### CRediT Author Statement: Rewati Prakash

Conceptualization, Writing-original draft, Methodology, Investigation, Formal analysis, Visualization. Benjamin N Forgie: Investigation, Formal Analysis; Alicia A Goyeneche: Supervision, Methodology; Edith Zorychta: Editing; **Abu Shadat M Noman**: Funding Acquisition; **Lucy Gilbert**: Funding Acquisition, Clinical Training; Carlos M. Telleria: Editing, Supervision, Resources, Project Administration, Funding Acquisition, Conceptualization.

## Acknowledgments

We thank Dr. Mark Carery of the Department of Pathology, University of British Columbia, for providing the patient-derived low-grade serous ovarian cancer cell models VOA6406, VOA1056, and VOA7681 generated in his laboratory. The results presented in this manuscript were previously presented, at least in part, in abstract form [56].

## Ethics approval and consent to participate

Not Applicable

## Figure Legends

**Supplementary Figure 1:**
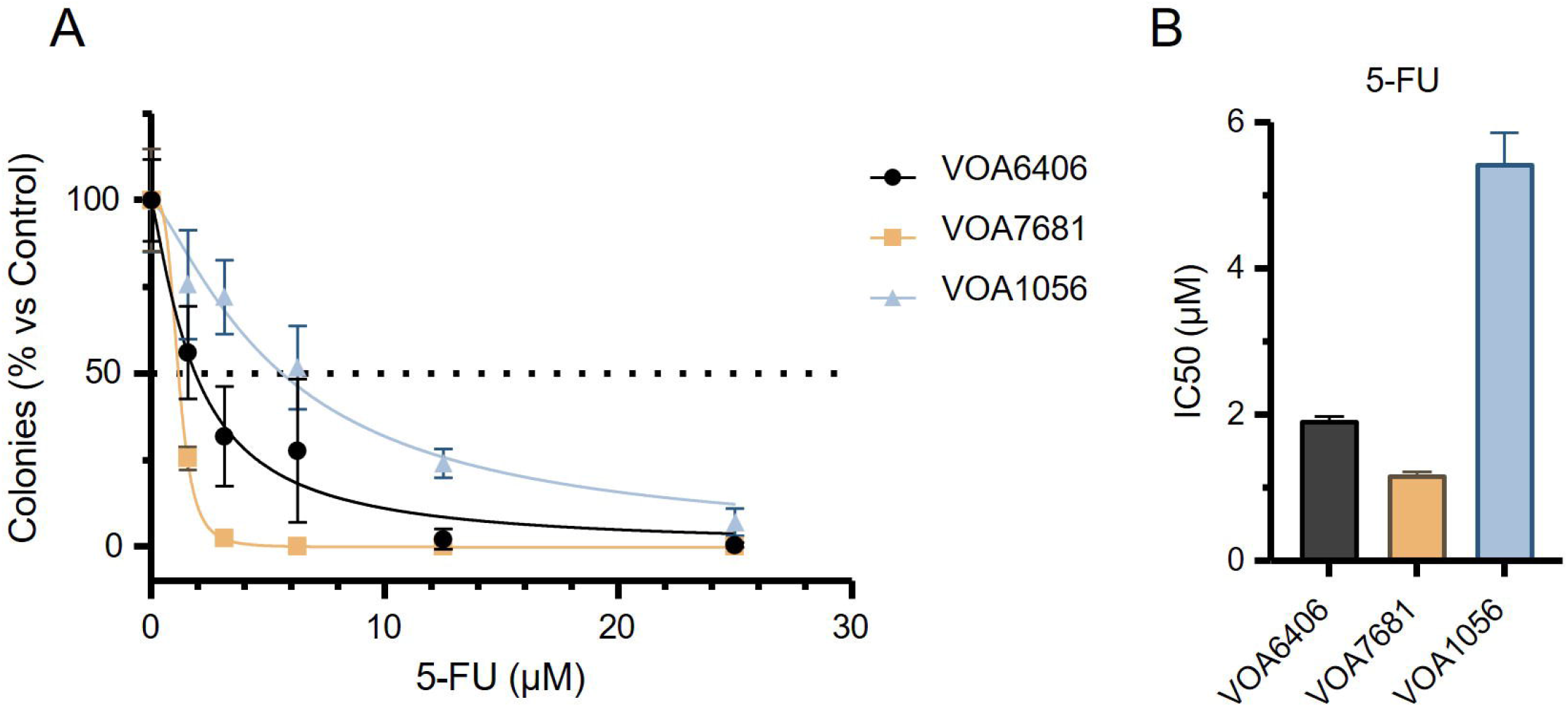
Clonogenic recovery dose-response of 5-FU-treated VOA6406, VOA7681, and VOA1056 cells (**A**). Half-maximal inhibitory concentration (IC50) of 5-FU clonogenic dose-response (**B**).

**Supplementary Table 1:** Details of Patient-Derived LGSOC cells.

| Cell Line | Tumor Details at Collection | LGSC Mutation Status | Treatment | References |
| --- | --- | --- | --- | --- |
| VOA6406 | Age: 56<br>Recurrent LGSOC | NRAS (Q61R;<br>c.182A>G)<br>WT: TP53, BRAF,<br>KRAS | Carboplatin-Paclitaxel | [46,57,58] |
| VOA1056 | Age: 62<br>Stage IIIC<br>Micropapillary serous<br>borderline ovarian tumor<br>(SBOT) with invasive<br>implants | NRAS (Q61R;<br>c.182A>G)<br>WT: TP53, BRAF,<br>KRAS | Naïve | [46,57-59] |
| VOA7681 | Age: 59<br>Stage IV | KRAS (G12V;<br>c.35G>T) | Naïve | [31,57] |
|  |  | WT: TP53, BRAF,<br>NRAS |  |  |

