## Supplementary material for "Oxaliplatin and 5-Fluorouracil induce p53-p21-pRb-associated cell cycle arrest and a transient senescence-like phenotype in patient-derived low-grade serous ovarian cancer cells": Uncropped Western Blot Images

**Original uncropped immunoblot images (Prakash et al.)**

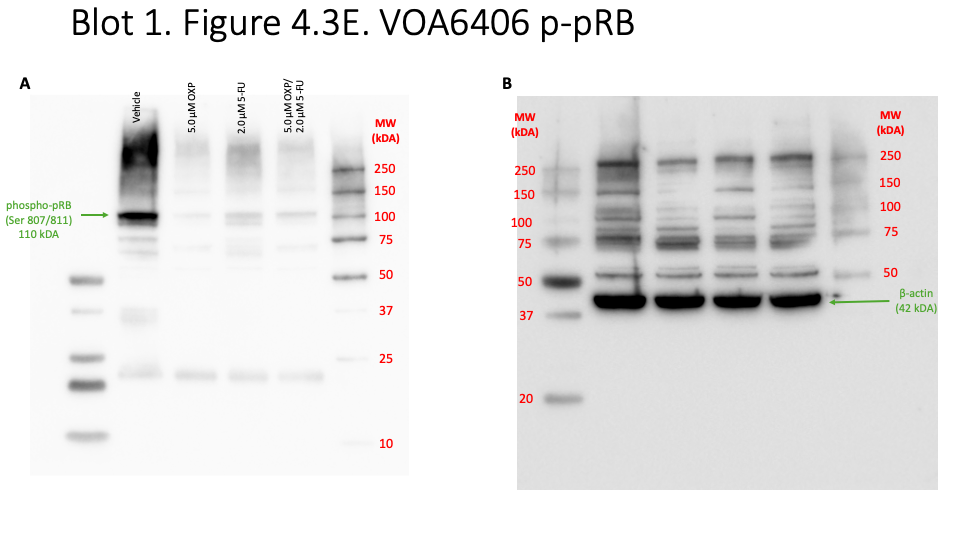
**Blot 1. Figure 2C. VOA6406 phosphorylated pRB**

VOA6406 cells were treated with 5.0 µM OXP and 2.0 µM 5-FU for 72 h. Western blot analysis was performed to measure specific protein expression. The blot was incubated overnight at 4 °C with anti-phospho-pRB (Ser 807/811) antibody (110 kDa) in 5% BSA-TBST (**A**), then with anti-β-actin antibody (42 kDa) (**B**).

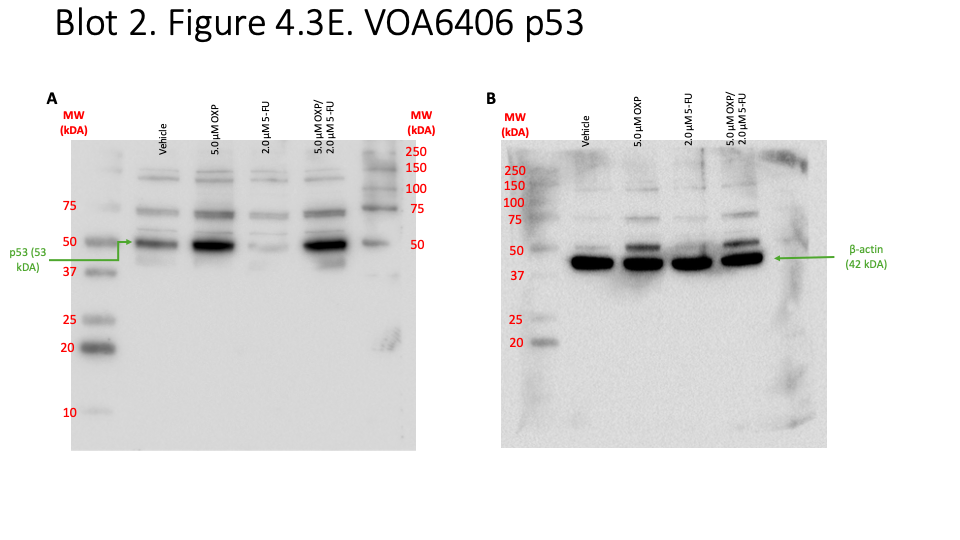
**Blot 2. Figure 2C. VOA6406 p53**

VOA6406 cells were treated with 5.0 µM OXP and 2.0 µM 5-FU for 72 h. We performed Western blot analysis to measure specific protein expression. The blot was incubated overnight at 4 °C with anti-p53 antibody (53 kDa) in 5% milk-TBST (**A**), then with anti-β-actin antibody (42 kDa) (**B**).

**Blot 3. Figure 2C. VOA6406 p21**

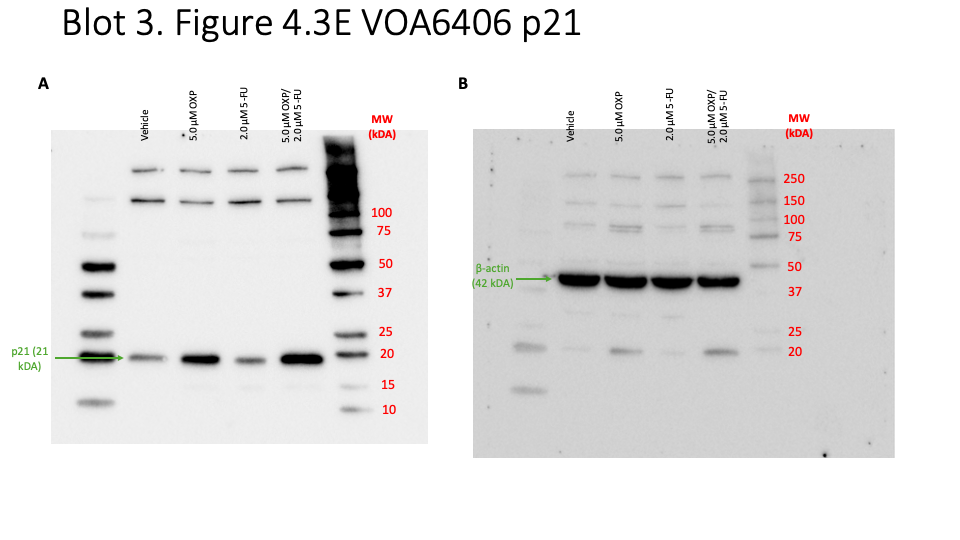
b

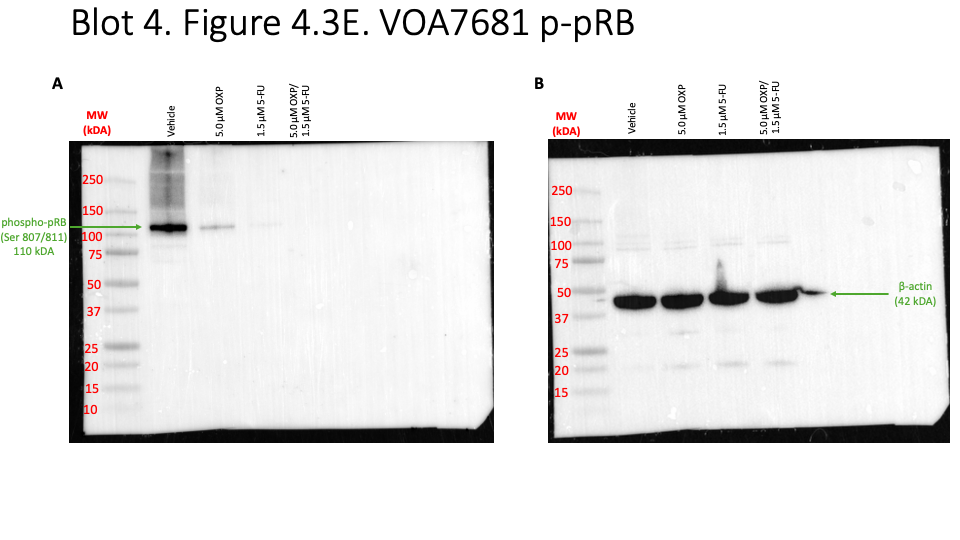
**Blot 4. Figure 2C. VOA7681 p-pRB**

VOA7681 cells were treated with 5.0 µM OXP and 1.5 µM 5-FU for 72 h. We performed Western blot analysis to measure specific protein expression. The blot was incubated with the primary antibody, anti-phospho-pRB antibody (110 kDA) overnight in 5% BSA-TBST at 4 °C (**A**), and then subsequently with anti-β-actin antibody (42 kDA) (**B**).

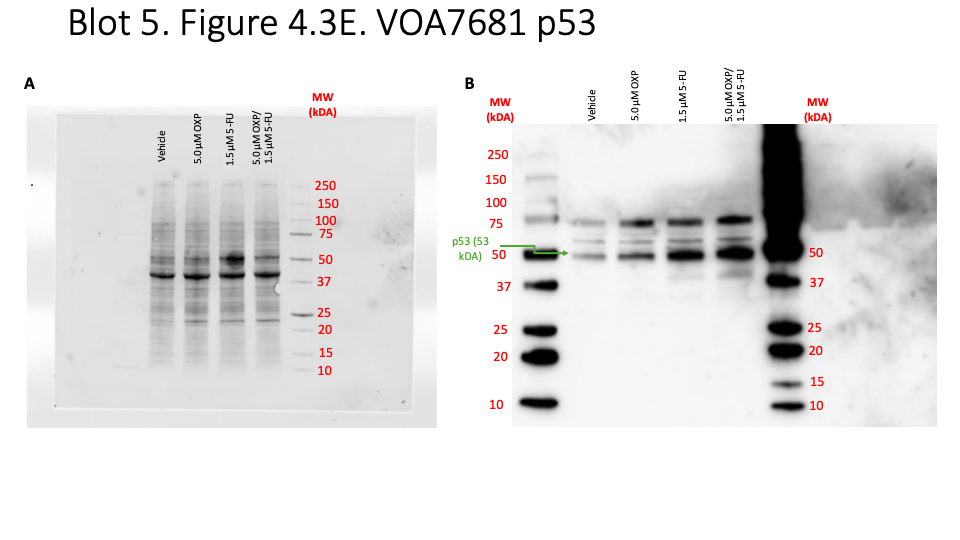

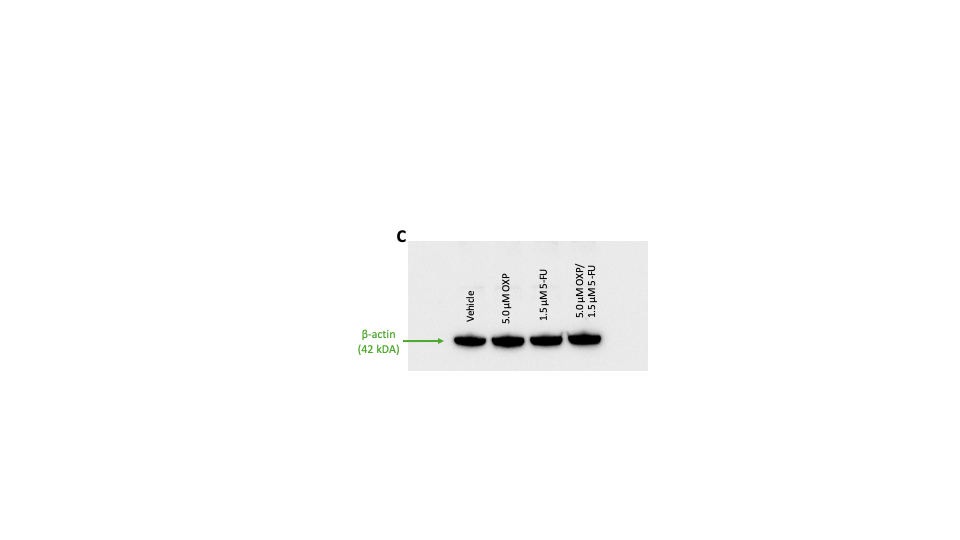
**Blot 5. Figure 2C. VOA7681 p53**

VOA7681 cells were treated with 5.0 µM OXP and 1.5 µM 5-FU for 72 h. Western blot analysis was performed to measure specific protein expression. The total protein load was visualized in the unstained membrane (**A**) before the blot was blocked and incubated with the primary antibody, anti-p53 antibody (53 kDA) overnight in 5% milk-TBST at 4 °C (**B**), and then subsequently with anti-β-actin antibody (42 kDA) (**C**).

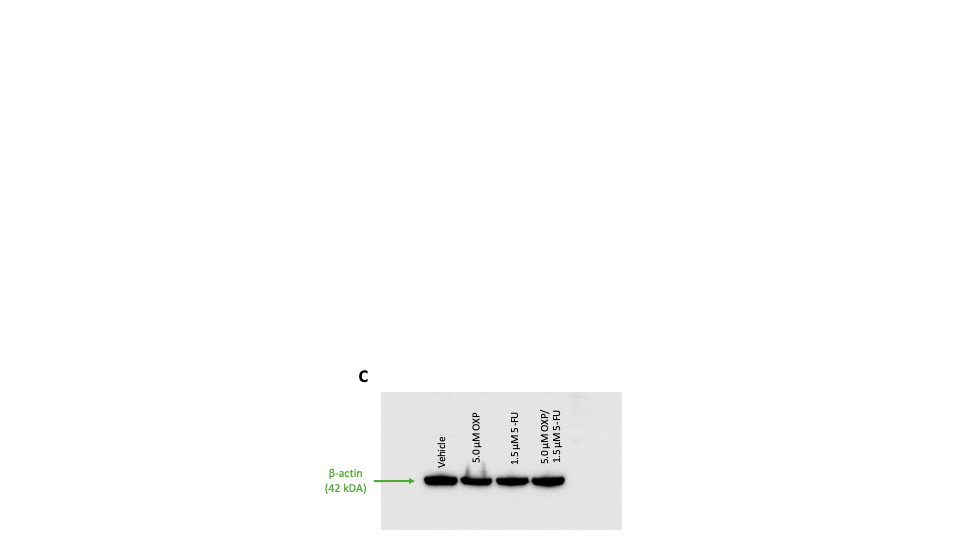

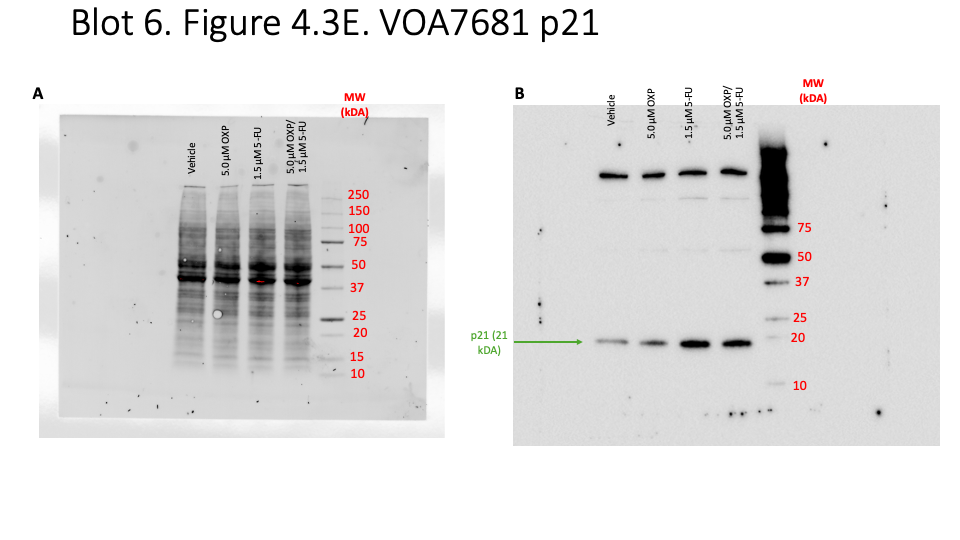
**Blot 6. Figure 2C. VOA7681 p21**

VOA7681 cells were treated with 5.0 µM OXP and 1.5 µM 5-FU for 72 h. Western blot analysis was performed to measure specific protein expression. The total protein load was visualized in the unstained membrane (**A**) before the blot was blocked and incubated with the primary antibody, anti-p21 antibody (p21 kDA) overnight in 5% milk-TBST at 4 °C (**B**), and then subsequently with anti-β-actin antibody (42 kDA) (**C**).

**Blot 7. Figure 3E. VOA7681 p16**

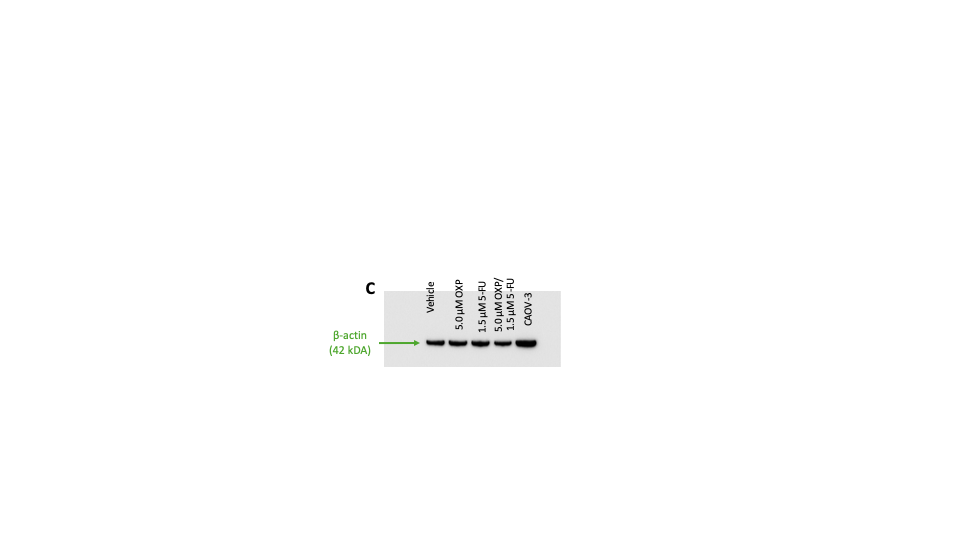

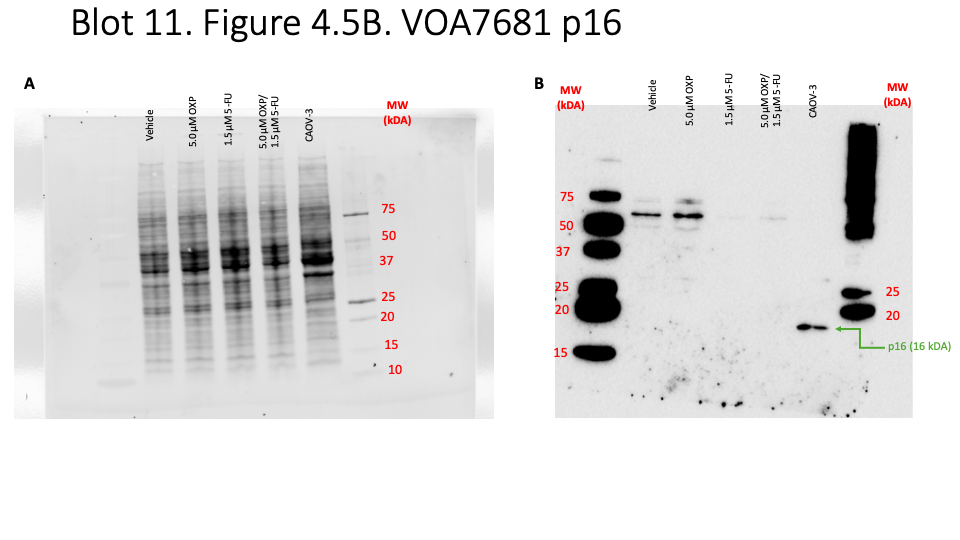

VOA7681 cells treated with 5.0 µM OXP and 2.0 µM 5-FU for 72 h, and untreated CAOV-3 cells were processed for western blot analysis to measure expression of specific proteins. The total protein load was visualized on the unstained membrane **(A)** before blocking and incubation with anti-p16 antibody (40 & 32 kDa) overnight in 5% milk-TBST at 4 °C **(B)**, followed by incubation with anti-β-actin antibody (42 kDa) (**C**).
